# Kmer-Node2Vec: a Fast and Efficient Method for Kmer Embedding from the Kmer Co-occurrence Graph, with Applications to DNA Sequences

**DOI:** 10.1101/2022.08.30.505832

**Authors:** Zhaochong Yu, Zihang Yang, Qingyang Lan, Yuchuan Wang, Feijuan Huang, Yuanzhe Cai

## Abstract

Learning low-dimensional continuous vector representation for short k-mers divided from long DNA sequences is key to DNA sequence modeling that can be utilized in many bioinformatics investigations, such as DNA sequence classification and retrieval. DNA2Vec is the most widely used method for DNA sequence embedding. However, it poorly scales to large data sets due to its extremely long training time in kmer embedding. In this paper, we propose a novel efficient graph-based kmer embedding method, named Kmer-Node2Vec, to tackle this concern. Our method converts the large DNA corpus into one kmer co-occurrence graph and extracts kmer relation on the graph by random walks to learn fast and high-quality kmer embedding. Extensive experiments show that our method is faster than DNA2Vec by 29 times for training on a 4GB data set, and on par with DNA2Vec in terms of task-specific accuracy of sequence retrieval and classification.

## I. INTRODUCTION

DNA/RNA sequence embedding technique has been used in many fields, such as pathogen detection [1], species traceability [2], microbiome analysis [3], etc. Recent DNA/RNA sequence embedding works [4], [5] first divide the long DNA sequence into kmer components, applies the skip-gram [6] to gain the kmer embedding for these kmer components, and then aggregates the kmer embedding to present the DNA sequence. Nevertheless, these methods scale poorly to large-scale data sets because they suffer huge time from training the kmer embedding. For example, DNA2Vec [4] needs **4.5 days** to train on 4GB DNA sequence data set. To tackle this concern, we propose a novel approach, Kmer-Node2Vec, which needs 3.7 hours (**29** times faster than DNA2Vec) with the better accuracy on the same data set.

Our idea comes from De Bruijn graph [7] (see Fig. 1(A)), which is applied for the genome assembly. In traditional sequencing algorithms [8] (e.g., Sanger sequencing or Next generation sequencing), reads (3mers) are represented as nodes in a graph, and edges represent alignments between the reads. Walking along a Hamiltonian cycle by following the edges in numerical order allows one to reconstruct the genome by combining alignments between successive reads. De Bruijn graph with applications to genetic sequences indicates that *graph structure* can be applied to represent genetic sequences well. Inspired by De Bruijn graph, we propose **the kmer co-occurrence graph** (see Fig. 1(B)) to describe the genetic sequences. Similar to De Bruijn graph, the kmers in a sequence (*3mers* in Fig. 1(B)) are represented as nodes in a kmer co-occurrence graph, and edges represent kmer co-occurrence relationship (alignments) between nodes, with edge direction as the kmer order. For example, the edge from GAA to AAG denotes that “AA” is the *co-occurrence 2mer* between G**AA** and **AA**G. However, different from De Bruijn graph, the number of co-occurrences of two kmers in the genetic sequence is used as the edge weight, and the kmer frequency in the sequence is used as the node weight, on the kmer co-occurrences graph.

**Fig. 1.**
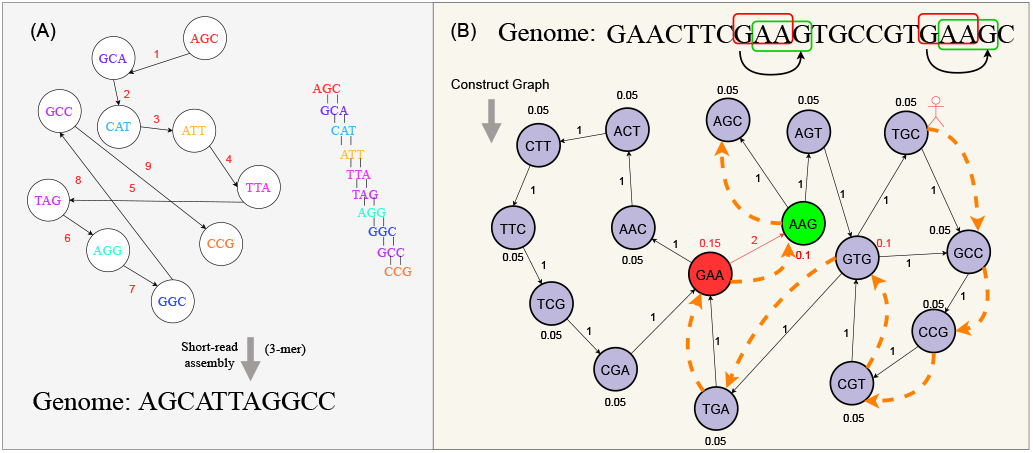
De Bruijn Graph vs. the Kmer Co-occurrence Graph. Fig. 1(A) describes De Bruijn graph for gene assembly. In traditional Sanger sequencing algorithms, reads (*3mers*) are represented as nodes in a graph, and edges represent alignments between reads. Travelling along a Hamiltonian cycle by following the edges in numerical order allows one to reconstruct the genome sequence by combining alignments between successive reads. Fig. 1(B) describes the 3mer co-occurrence graph. This 3mer co-occurrence graph is built from the DNA sequence “GAACTTCGAAGTGCCGTGAAGC”. An edge is directed from 3mer node *v* to node *x* if they overlap by two continuous nucleotides with node *v*’s suffix being the node *x*’s prefix, i.e., from “GAA” (the red node in genome) to “AAG” (the green node), and “AA” as their overlap. The pair of “GAA” to “AAG” occurs *twice* in the sequence, so that the weight of the graph edge from “GAA” to “AAG” is 2. The dashed line with an arrow describes a path of a random walk with “TGC” as the starting node and “AGC” as the ending node.

Our method, Kmer-Node2Vec^1^, samples kmers by random walk traveling on the kmer co-occurrence graph, and learns kmer embedding from these kmers. First, Kmer-Node2Vec uses kmer co-occurrence information in DNA sequences to construct a kmer co-occurrence graph. Second, it performs fast and efficient random walk traveling on the graph to sample kmer sequences. Third, Skip-Gram [6], a language embedding model, is applied to these sampled kmer sequences to learn the kmer embedding. We argue that training the kmer embedding on a kmer co-occurrence graph is effective and efficient for the following reasons.

i. *Effectiveness*. With random travelling on the kmer co-occurrence graph, our method is able to capture both **local** (in a local sliding window) and **global** (among different sequences) kmer co-occurrence information together, and use this information to embed kmers under the assumption that highly co-occurring kmers should have similar meaning [4].
ii. *Efficiency*. Two points motivate the improvement of our method’s efficiency. First, DNA2Vec requires a long period to train the kmer embedding, since **its** training corpus of kmers is huge. DNA molecule in the nucleoid of Take an Escherichia coli, whose DNA sequence is 4.7 million bp, as an example. This bacterial genome will be fragmented into 4.7 million kmers in kmer representation, which indicates that even a E coli’s data size is equivalent to a 4.7 million words long novel (one kmer corresponds one word in word2vec [6]). That is equal to **four** times of total words in “Gone with the Wind”. Second, another interesting observation on DNA sequences is that there are a large number of **same** genetic segments among various species. For example, comprehensive analysis of chimpanzee and human chromosomes reveals average DNA similarity of 70% [9]. Meanwhile, SARS-CoV-2 shares 79% genome sequence identity with SARS-CoV and 50% with MERS-CoV [10]. DNA2Vec wastes too much computation by sliding a window on these same genetic segments. Based on this observation, we emphasize that the kmer co-occurrence graph, which has already captured all the **statistics** of genetic sequences, is a better model to analyze the genetic sequences. The number of nodes in the kmer co-occurrence graph is not large since the maximum number of graph nodes is 4^*k*^ (e.g., 8-mer graph contains 4^8^ (65536) nodes), and the kmer co-occurrence graph is highly sparse because the maximum edges for each node are four as there are only four kinds of base-pair (A, T, C, G) (e.g., in 8mer co-occurrence graph, the density is no more than 0.01%, 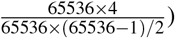). Meanwhile, these above-mentioned same DNA segments only change the edge weights of the kmer co-occurrence graph, but the size of nodes and graph density do not have any alteration. Thus, it does not affect the Kmer-Node2Vec’s performance.

## II. Kmer-Node2Vec Algorithm

Fig. 2 clearly shows our Kmer-Node2Vec’s framework, which includes constructing the kmer co-occurrence graph, randomly travelling on the graph, learning kmer embedding by Skip-Gram [9], calculating the sequence embedding, and applying sequence embedding for bioinformatics tasks.

**Fig. 2.**
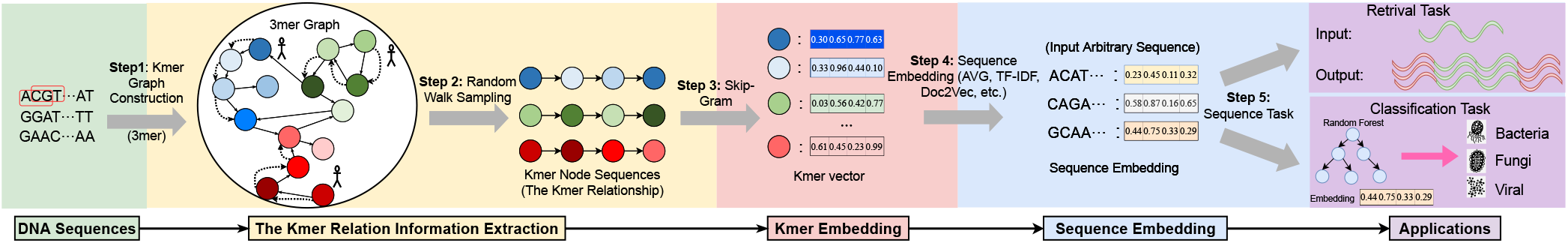
The overall framework of Kmer-Node2Vec algorithm for kmer embedding. In step 1, Kmer-Node2Vec takes DNA sequences as input and constructs a kmer co-occurrence graph based on them. In step 2, Kmer-Node2Vec uses random walks to extract kmer relation sequences. In step 3, Kmer-Node2Vec feeds these sequences to Skip-Gram [6] and produces a kmer vector for each kmer (kmer embedding). In step 4, these kmer vectors can be used to compute the DNA sequence embedding using AVG, TF-IDF, Doc2Vec [11]. In step 5, we apply the sequence embedding for bioinformatics tasks, such as sequence classification, sequence retrieval and so on.

### 1) The Kmer Co-occurrence Graph Construction

We map *numerous* DNA sequences to a kmer co-occurrence graph that corresponds to a weighted directed graph whose nodes represent unique kmers, whose edges represent cooccurrences between kmers with edge direction denoting the order of kmers. To simply illustrate, we use only one DNA sequence to construct the graph. Let a sequence be *GAACTTCGAAGTGCCGTGAAGC* and *k* = 3, the sequence is represented as a series of *3mers* = {*GAA, AAC, ACT, CTT, TTC, TCG, CGA, GAA, AAG, AGT, GTG, TGC, GCC, CCG, CGT, GTG, TGA, GAA, AAG, AGC*}. Then we take each unique 3mer as a node of the graph. An edge is drawn between a kmer and its following kmer. Fig. 1(B) corresponds to the graph that we construct.

#### Weight on edges

The co-occurrence number of two kmers in the sequences is used as their edge weight. A weighted adjacency matrix *W* is used to store the edge weights, and *W*_*vx*_ is the edge weight from node *v* to node *x*.

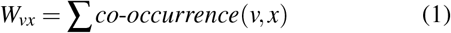

where *co*-*occurrence*(*v, x*) is the number of times that the kmer *v, x* appear together from left to right. For instance, the number of occurrences of *GGA* ⇒ *GAA* is 2, thus *W*_*GGA,GAA*_ is set as 2 (see Fig. 1(B)).

#### Weight on nodes

The node weight is used to determine the sampling times of random walks. We take probability *PW*_*v*_ to indicate how kmer *v important* is in the DNA corpus, that is, the node weight in the graph. Intuitively, the kmer frequency can be applied to measure the importance score.

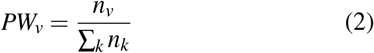

where *PW*_*v*_ is the important score for kmer *v, n*_*v*_ denotes the occurrence number of the kmer *v*, and ∑_*k*_ *n*_*k*_ represents the occurrence number of all kmers. Taking Fig. 1(B) as an example, ***GGA*** appears 3 times and the occurrence number of total kmers is 20, therefore 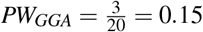.

### 2) Kmer Node Sampling

This random walk sampling process [12] iteratively explores the local structure of the kmer co-occurrence graph to estimate the **proximity** between kmer nodes. Generate a random walk from the current node, and select the random neighbors of the current node as a candidate based on the transition probability on the kmer co-occurrence graph. Take a hypothetical case in Fig. 1(B) as an example to explain how we use our sampling strategy to sample a sequence (nodes) from the kmer co-occurrence graph. We select *TGC* as the starting node to execute a random walk of walk-length 8; the random walk starts from *TGC*, passes through *GCC, CCG*, …, and finally arrives at *AGC*. The sample of the sequence (nodes) obtained by this random walk is exactly *TGCCGTGAAGC* = *TGC* → *GCC* → *CCG* → *CGT* → *GTG* → *TGA* → *GAA* → *AAG* → *AGC*. We iterate over all the graph nodes to execute numerous random walks to sample kmer sequences. To *better estimate* the kmer proximity, we carefully design the random walk procedure with the following *three* characteristics.

First, the transition probability is affected by directed edge weights. This forces random walks to preserve the kmer cooccurrence information as much as possible. The probability of traversing from node *v* to its neighbour node *x* is *W*_*vx*_ (see Eq. 1) divided by the sum of all neighbouring *directed* weights. As shown in Fig. 1, probability to traverse from node **GAA** to node **AAG** is 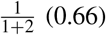 (0.66).

Second, each starting node should execute sufficient random walks to obtain a qualified sample for kmer embedding training. A node’s number of walks should be in line with its role in the graph, as important nodes should enjoy more random walks than others. We assign each node with a node weight (see Eq. 2) that reflects a node’s importance. The number of random walks rooted at node *v* is computed using Eq. 3:

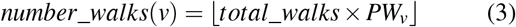

where *total*_*walks* is the total number of random walks set by the user. For example, in Fig. 1(B), let *total*_*walk* = 200, *number*_*walks*(*GAA*) should be 200 × 0.15 = 30. That means there are 30 sampled sequences that start from node *GAA*.

Third, to capture both homophily and structural proximity between kmer nodes of the graph, we employ Node2Vec [12] second-order sampling strategy in Kmer-Node2Vec. In addition, Pecanpy [13], which offers a parallelized and accelerated Node2Vec in python, is applied in our experiments.

### 3) Training Kmer Embedding with Skip-Gram

We use Skip-Gram to train the kmer embedding on the sampling corpus. The sampling corpus is composed of numerous random walks, which can be thought of kmer node sequences. Skip-Gram maximizes the *co-occurrence probability* among the kmers that appear within a local context in a kmer node sequence. In that way, Skip-Gram will produce very similar vector representations for two kmers if they are *frequently* within a local context of sampled kmer node sequences. With our sampling strategy, there is a **high** likelihood that such two kmers are also *frequently* within a local context in DNA sequences, and thus have similar meaning [4].

### 4) Theoretical Analysis on Performance

We pay attention to the runtime gap between Kmer-Node2Vec and DNA2Vec. Denoting *N* as the total corpus size and *V* as the uniquekmers vocabulary count, DNA2Vec has a high time complexity in training kmer embedding with Skip-Gram, which is *O*(*Nlog*(*V*)). *N* will be very big on genetic data sets (e.g., *N* ≈ 1.1e9 on 1GB data set, and ≈ 4.5e9 on 4GB data set). However, in Kmer-Node2Vec, the training corpus for Skip-Gram is our sampling corpus whose size is *nl*, where *n* is the number of random walks and *l* is the walk length. The time complexity is reduced to *O*(*nllog*(*V*)), since *nl* is smaller than *N* (*nl* ≈ 1.9e8 for both 1GB and 4GB data set). On large-scale data sets, *V* is the fixed value, and *nl* is also set by the user so that our method’s training time is relative **stable** for various-size data sets.

## III. Experiment

### A. Experimental Settings

We use various data sets in our experiments (see Table I), and set
kmer size as 8 to train the model.

**TABLE I.** The statistic of data sets

| Data set | Total size | Total sequence number | Average sequence length |
| --- | --- | --- | --- |
| Fungi | 142MB | 2,007 | 71,670 bp |
| Virus | 380MB | 13,274 | 29,563 bp |
| Bacteria | 706MB | 28,115 | 25,858 bp |
| 1G data set | 1GB | 43,145 | 26,644 bp |
| 2G data set | 2GB | 65,614 | 28,947 bp |
| 3G data set | 3GB | 95,991 | 30,599 bp |
| 4G data set | 4GB | 129,999 | 34,247 bp |

### B. Efficiency of Kmer-Node2Vec algorithm

Fig. 3(a) and Fig. 3(b) summarize the runtime and memory usage for DNA2Vec [4]. The empirical results show that: (i) Kmer-Node2Vec is much faster than DNA2Vec in training speed on **all** data sets, with the former 27 times faster than the latter, on average; (ii) As training DNA data grows, Kmer-Node2Vec’s runtime increases slower than DNA2Vec’s, indicating that larger data will see a greater runtime gap between them. For example, from 1G data set to 4G data set, Kmer-Node2Vec’s runtime increases slightly from 80min to 230min, whereas DNA2Vec’s runtime rises from 1,780min to 6,600min; (iii) Kmer-Node2Vec consumes more memory usage than DNA2Vec, because the kmer co-occurrence graph needs more memory to storage. Fortunately, the memory usage of our method intends to remain unchanged once reaching the peak (10GB), since the size of the kmer co-occurrence graph and sampled sequences can be viewed as constant.

**Fig. 3.**
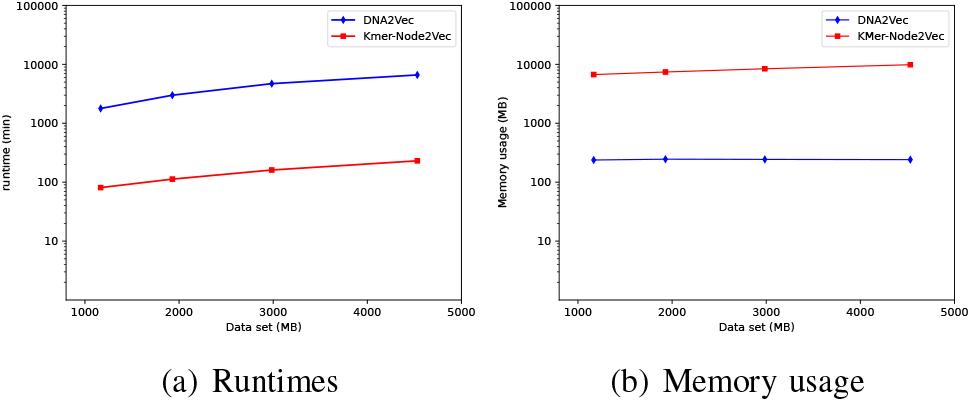
Runtimes and Memory usage on four data sets (1G-4G). Note that the ordinate scale is arranged in logarithmic base 10.

### C. Accuracy Evaluation

#### 1) Retrieval Task

This task is to evaluate the accuracy with a baseline approach, **BLAST+** [14]. The DNA sequence *retrieval system* using either our method or DNA2Vec is described here: (i) DNA sequences in *three* data sets of Fungi, Virus and Bacteria (see Table I) are split into non-overlapping segments of 150bp, denoted as *S*_150*bp*_; (ii) these segments are transformed to vectors to load into Faiss [15] system. HNSW [16] index is built for vector retrieval in the Faiss system; (iii) given a query sequence, we use its sequence vector (an average of learned kmer vectors) to search in the Faiss to find Top-*K* segment vectors that are highly similar to the query sequence vector. All parameters for BLAST+ and Faiss in the experiments are adjusted to achieve the best performance.

Specifically, we randomly extract 1,000 DNA segment sequences (15bp-75bp) from *S*_150*bp*_ as query sequences and take the corresponding labels as the ground-truth. We feed each query sequence into the retrieval system and Top-*K* similar DNA sequence segments, denoted as *S*_*K*_, are returned. Top-*K* accuracy is used to measure the proportion of the correct label is within the Top-*K* sequences returned by the retrieval system, denoted as 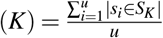, where *K* denotes the number of query’s nearest neighbors to be found, *u* represents the number of queries, and *s*_*i*_ is the correct segment label as the ground-truth of the query.

Overall, in Fig 4, Top-*K* accuracy of Faiss is slightly lower than BLAST+, but Faiss takes a much shorter retrieval time than BLAST+. For example, Faiss is about **1.4 times** and **3.3 times** faster than BLAST+ using the minimum data set (Fungi) and the maximum one (Bacteria), respectively. Meanwhile, Kmer-Node2Vec’s Top-*K* accuracy is higher than DNA2Vec’s, since our method can capture both local and global co-occurrences between kmers. Because Faiss with Kmer-Node2Vec typically achieves high search accuracy of **99.5%** at the Top-20 level, it is adequate to satisfy the real application.

**Fig. 4.**
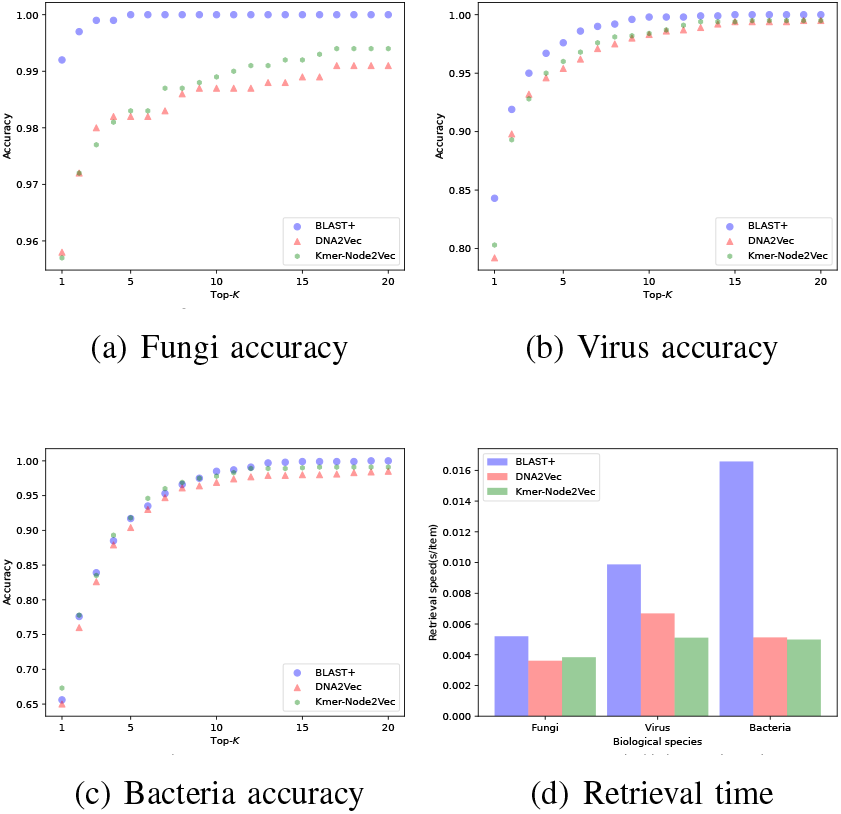
Fig 4(a), (b), (c) compare the sequence retrieval accuracy. Fig 4(d) shows the retrieval time.

#### 2) Classification Task

This task first randomly chooses 18,000 DNA sequences of three species (6,000 for fungi, virus and bacteria, respectively). We apply either our method or DNA2Vec to calculate the sequence embedding (an average of learned kmer vectors), and build the random forest classifier with the sequence embedding as features to classify these sequences into their corresponding species (fungi, viral, bacteria). Precision, Recall and F1-score [17] computed using 10-fold cross-validation are used to test the performance of classification. Experimental results show that the precision, recall and F1-score for our method and DNA2Vec are (0.942, 0.860, 0.896) and (0.936, 0.849, 0.887), respectively, indicating that our method outperforms DNA2Vec (**1.01%** improvement for F1-score) on the classification task.

## Footnotes

1 Kmer-Node2Vec’s source code can be downloaded from https://github.com/caiyuanzhe/kmer-node2vec

